# Supramodal Sentence Processing in the Human Brain: Fmri Evidence for the Influence of Syntactic Complexity in More Than 200 Participants

**DOI:** 10.1101/576769

**Authors:** Julia Uddén, Annika Hultén, Jan-Mathijs Schoffelen, Nietzsche Lam, Karin Harbusch, Antal van den Bosch, Gerard Kempen, Karl Magnus Petersson, Peter Hagoort

**Affiliations:** Max Planck Institute for Psycholinguistics, Nijmegen, the Netherlands; Donders Institute for Brain, Cognition and Behaviour, Centre for Cognitive Neuroimaging, Radboud University, the Netherlands; Department of Computer Science, University of Koblenz-Landau, Koblenz, Germany; Department of Linguistic, Stockholm University, Sweden; Department of Psychology, Stockholm University, Sweden.

**Keywords:** complexity, fMRI, sentence processing, supramodal, unification

## Abstract

This study investigated two questions. One is to which degree sentence processing beyond single words is independent of the input modality (speech vs. reading). The second question is which parts of the network recruited by both modalities is sensitive to syntactic complexity. These questions were investigated by having more than 200 participants read or listen to well-formed sentences or series of unconnected words. A largely left-hemisphere fronto-temporoparietal network was found to be supramodal in nature, i.e. independent of input modality. In addition, the left inferior frontal gyrus (LIFG) and the left posterior middle temporal gyrus (LpMTG) were most clearly associated with left-branching complexity. The left anterior middle temporal gyrus (LaMTG) showed the greatest sensitivity to sentences that differed in right-branching complexity. Moreover, activity in LIFG and LpMTG increased from sentence onset to end, in parallel with an increase of the left-branching complexity. While LIFG, bilateral anterior and posterior MTG and left inferior parietal lobe (LIPL) all contribute to the supramodal unification processes, the results suggest that these regions differ in their respective contributions to syntactic complexity related processing. The consequences of these findings for neurobiological models of language processing are discussed.

## 1. INTRODUCTION

In order to extract meaning from the orthographic patterns or from the speech sounds, multiple processing steps are involved. One important step is to retrieve relevant word information from long-term memory (the mental lexicon). This information includes the morphological make-up of words, their syntactic features, and lexical aspects of their meaning. But this is not enough. In many cases a simple concatenation of individual word meanings will not result in a correct interpretation (Jackendoff 2002). The reason is that in language, words that belong together often do not go together (Lashley 1951). This is what linguists refer to as non-adjacent dependencies between the lexical elements that make up an utterance. How to combine word information retrieved from memory into representations of sentence-level meaning that are constructed on the fly is what we refer to as unification (Hagoort, 2005, 2013, 2014; Vosse & Kempen, 2000, 2009). The number of non-adjacent elements that have to be kept on-line determines unification complexity. In this large fMRI study on sentence processing (N=204), we address two outstanding questions: (i) to what extent is the network subserving unification operations independent of the modality of input (spoken and written)? This was investigated by confronting half of the participant with the materials in spoken format and half with the same materials in written format; (ii) which nodes in the language network are modulated by variation in syntactic complexity. For each sentence presented to the participant, we calculated a measure of complexity, which allowed us to identify the areas that were most sensitive to complexity variations.

### Modality-independence of Unification

It is generally assumed that structure building processes in the spoken and written modalities are subserved by similar modality-independent operations. For instance, in the processing model of structure building called the *Unification framework* (Vosse and Kempen 2000, 2009), the attachment of each new lexical item to the incrementally constructed syntactic representation of the sentence, is identical for visual and auditory language input, but this has not been explicitly tested. In the Memory, Unification and Control framework (Hagoort 2005, 2013), the word information mainly stored in the temporal lobe includes specifications of syntax, morphology and information about word meaning (Joshi and Schabes 1997; Vosse and Kempen 2000). The process of unifying these lexical information types with the sentence and discourse context is constrained by lexical features in a process assumed to be supramodal. It has been suggested, that after a forward sweep from sensory cortex to the left temporal and parietal lobe, top-down signals from the left inferior frontal gyrus (LIFG) re-enter the posterior regions in cycles of reactivation (Baggio and Hagoort 2011), establishing a unification network with the involvement of at least two left hemisphere regions (left temporal/parietal cortex and LIFG, where the LIFG is the higher level node in the network). Visual and auditory language processing streams are hypothesized to converge on this supramodal unification network during comprehension. The unification network thus includes both a frontal and a temporo-parietal node, but the LIFG is thought to be crucial for the higher level unification processes whereas the mental lexicon (or Memory component) is thought to reside in the temporo-parietal node.

### Sentence complexity

The second aspect that we addressed is related to sentence complexity. Processing complexity in language processing is often due to the fact that words that belong together do not always go together; that is, they do not appear in adjacent positions. In a recent study (Futrell et al., 2015) it was found that there is an almost universal tendency (based on an analysis of 37 languages) to disprefer sentences in which structurally related words are far apart, presumably as a result of the extra processing costs associated with non-adjacency. Nevertheless, non-adjacency is a common phenomenon in language processing and a hallmark of human languages. It occurs not only in sentences with a left-branching structure, but also in sentences with right-branching structure. There is evidence that on the whole left-branching structures are harder to process than right-branching structures. An increased cost of maintenance and structure building for left-branching sentence aspects compared to right-branching sentence aspects was first suggested by Fodor and colleagues (Fodor et al. 1974). This claim of an added processing load for left branching structures has been supported by evidence from both production and comprehension studies (Kemper 1986, 1987; Cheung and Kemper 1992; Norman et al. 1992).

Snijders et al. (2009) identified the LIFG and the MTG as core unification regions in a study that contrasted sentence processing with the processing of word-lists. In the current study, we used a similar paradigm, but extended it with an explicit manipulation and measure of sentence complexity. The neuroimaging literature on sentence complexity is substantial (among many others, see Caplan et al. 1998; Cooke et al. 2002; Peelle et al. 2004; Makuuchi et al. 2009; Meltzer et al. 2010; Santi and Grodzinsky 2010), but most often the studies have been restricted to (a) comparing two conditions of complex vs simpler sentences and (b) to measuring one sensory modality only (but see Michael et al. 2001; Constable et al. 2004; Shankweiler et al. 2008; Braze et al. 2011). When comparing two conditions of sentence complexity, most studies have realized this manipulation by comparing object-relative to subject-relative sentences, the former ones known to be more complex than the latter ones (e.g. Michael et al., 2001; Cooke et al., 2002; Constable et al., 2004; Peele et al, 2004.; Meltzer et al., 2010; Braze et al., 2011). In addition, some studies (e.g. Stromswold, 1996; Caplan et al., 1998 and 2000; Makuuchi et al., 2009; Santi and Grodzinsky, 2010) compared more complex center embedded sentences to simpler right branching sentences. In our study, a complexity measure is instead used to calculate the processing complexity for each individual sentence.

Our complexity measure is motivated by two observations, first: (a) the finding that sentences with a left-branching structure (see Figure 1) are particularly hard to process (Kemper 1986, 1987, Cheung 1992, Norman 1992). We thus separated left-branching from right-branching complexity. The second motivating observation (b) is that there is a high processing cost related to building sentence structure with multiple simultaneous non-local dependencies This is found for both for natural (Makuuchi et al., 2009) and artificial grammars (Bahlmann et al. 2008; Makuuchi *et al.* 2009; de Vries et al. 2012; Udden and Bahlmann 2012; Udden et al. 2012). Based on these two observations, our measure quantifies the amount of simultaneous left-branching non-local dependencies in a sentence (see Figure 1). It is of central importance that the sentence complexity is related to the incrementality of parsing, from left to right.

**Figure 1.**
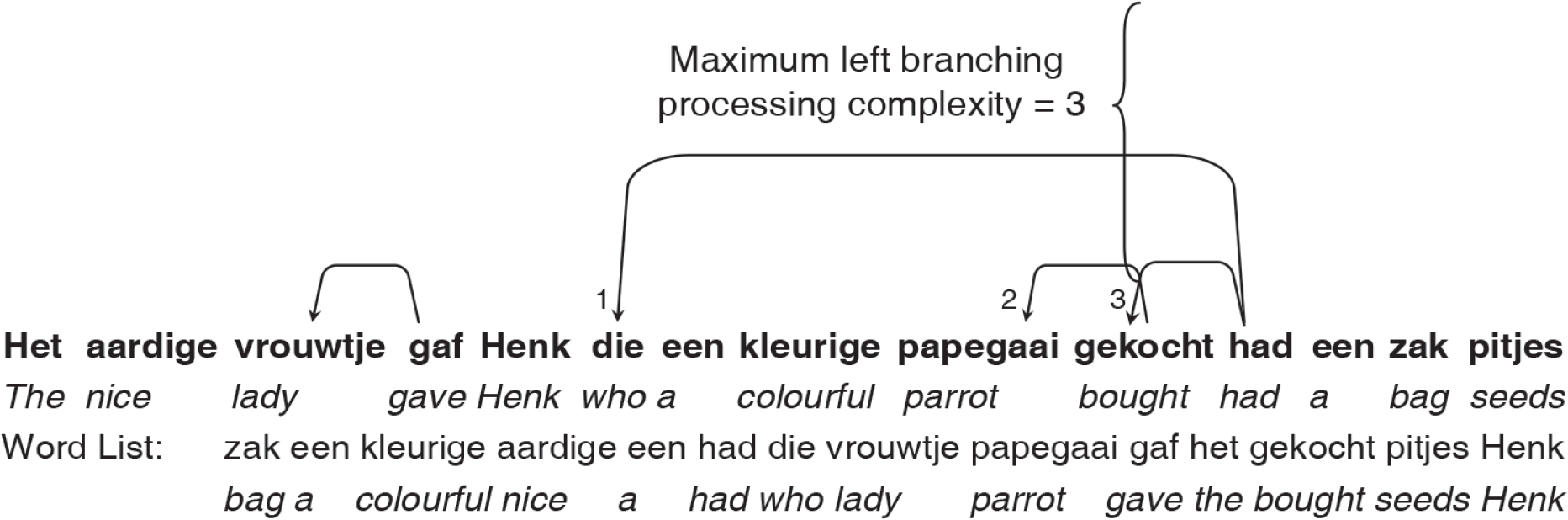
Left-branching dependencies point leftward from head to dependent. The left-branching processing complexity measure is calculated per sentence, as the maximum simultaneous non-resolved left-branching verbal dependencies (i.e. maximum number of dependents not yet assigned to a verb during the incremental parse). For the example sentence, this number equals 3 (reached after retrieval of the word ‘gekocht’). One of the open dependencies (from ‘papegaai’) is resolved after reading ‘gekocht’ and two other at ‘had’, since the participle (‘gekocht’) needs to be bound to an auxiliary (‘had’), and the auxiliary needs to fill its subject argument with an antecedent that has the right number marking (i.c., ‘Henk’, singular). Included below the sentence are: a literal translation to English, together with the corresponding word-list with translation.

To illustrate the importance of left to right processing, the reader might be helped by thinking of a *stack*, storing words that cannot yet be unified with the rest of the sentence structure. Each time a left-branching dependency is open, an element is pushed on the stack, and only when it is closed is the element popped. The left-branching complexity measure thus corresponds to the stack depth used for a sentence. However, we do not want to make a strong assumption that there is a buffer (in the form of a stack or otherwise). If no buffering of words occurs during incremental processing, the complexity measure still singles out the sentences which have a high unification load, for the following reason: multiple simultaneously open dependencies lead to more options for the dependent to be unified with a head, when the head arrives. The number of simultaneously open dependencies is thus a complexity measure probing relevant syntactic processes independent of the existence of a stack or other buffers.

The left-branching complexity was of greatest interest in our neuroimaging study since we predicted that it would be associated with the greatest processing difficulties. Our left-branching complexity measure quantifies the maximum simultaneous number of dependents not yet assigned to the head of a *verb phrase*. Unification happens between the head of a phrase and its arguments. For instance, in a noun phrase the determiner and the adjective need to be unified with the noun as the head of the noun phrase (e.g., “The nice lady”; see Figure 1). The longer the distance between the heads and their arguments, and the more dependents and arguments there are to be maintained, the larger the processing demands (see Figure 1). In this case, verb phrases are especially relevant, since in general the verb is the nucleus of the proposition that is expressed in the sentence. For instance, upon reading the verb *kick*, a noun or the name of a person is expected to fill the argument slot for the agent of the action specified by the verb (e.g., *the man*). In addition, another noun is expected to take the argument slot for the undergoer of the action (e.g., *the ball*, as in “the man kicks the ball”). We expected processing to be more demanding if the arguments precede the head, since the head is a stronger predictor for following arguments than the arguments are predictors for a following head. Therefore, we only count dependencies with a verbal head when we calculate the processing complexity.

To increase the sensitivity for unification associated with sentence complexity, we made sure that there was enough variance in the sentence structures. This was achieved by introducing relative clauses in half of our sentence material, while the other half had mixed sentence structures without relative clauses. Left-branching complexity was higher on average for sentences with relative clauses (see the methods section 2.2). The analysis was however not a standard analysis comparing two complexity conditions, but a parametric analysis probing for the sentence complexity effect across all sentences in the experiment.

In addition to localizing a supramodal structure building network in the brain, we also wanted to characterize the temporal dynamics of this network over the course of the sentence. For this purpose, we divided the sentences into four time-bins of equal length, and formed linear contrasts to test increases and decreases in activity over the course of a sentence. For this analysis, we focused on predefined regions of interest (ROI) found to be involved in structural unification (Snijders et al., 2009). Based on a recent meta-analysis of fMRI-studies of both syntactic and semantic sentence-level unification, the LIFG and the left posterior middle temporal gyrus (LpMTG) were chosen as primary regions of interest (Hagoort and Indefrey 2014). In the light of current debates on the role of left anterior temporal lobe in structure building processes during sentence processing (but see Pallier et al. 2011; Brennan et al. 2012), we also included a third ROI, based on coordinates from the study by Brennan et al.(2012), in a region in the left anterior middle temporal gyrus (LaMTG).

## 2. MATERIALS AND METHODS

### 2.1 Participants

A total of 242 participants volunteered to participate in a larger study – the MOUS study (Mother of all Unification Studies), where all participants took part in an fMRI and an MEG (magnetoencephalography) session. Of these, 38 participants were excluded. A full list of reasons for exclusion can be found in the supplementary material. The resulting 204 native Dutch speakers had a mean age of 22 years (range: 18 - 33 years). Half of the participants read sentences and word-lists (visual group, 102 participants, 51 men), and the other half listened to auditory versions of the same materials (auditory group, 102 participants, 51 men). The study was approved by the local ethics committee (CMO, the local “Committee on Research Involving Human Subjects” in the Arnhem-Nijmegen region) and followed guidelines of the Helsinki declaration.

### 2.2 Language stimuli

The stimuli consisted of 360 sentences and their 360 word-list counterparts. Sentences were constructed to vary in complexity. One way to make complexity vary is by introducing a relative clause (Gibson 1998). Half of the sentences contained a relative clause, the other half of the sentences were without a relative clause. Table 1 presents examples of the materials. The sentences varied between 9 and 15 words in length.

**Table 1.** Exemplar sentences and word-lists in Dutch, and literal English translation.

| Sentence | Word-list |
| --- | --- |
| Het aardige vrouwtje gaf Henk die een kleurige papegaai gekocht had een zak pitjes.<br>The nice lady gave Henk, who had bought a colorful parrot, a bag of seeds. | zak een kleurige aardige een had die vrouwtje papegaai gaf het gekocht pitjes<br>bag a colorful nice a had who lady parrot gave the bought seeds |
| Dit zijn geen regionale problemen zoals die op de Antillen.<br>These are not regional problems such as those on the Antilles. | zoals geen die Antillen problemen regionale zijn de dit op<br>such as not those Antilles problems regional are the these on |

For each sentence, a corresponding word-list was created by scrambling the words in the sentence so that three or more consecutive words did not form a coherent fragment. Since the sentence-and word-list conditions contain the same words, the comparison of sentences with word-lists allowed us to probe the sentence-level unification process, while controlling for the retrieval of lexical items. We calculated the complexity of each sentence on the basis of the dependency structure, separately for left-and right-branching sentence aspects. For the left-branching processing complexity measure, at each word, we calculated the number of dependents that had not yet been attached to a verbal head (i.e., the head had not yet been encountered). The maximum number over the entire sentence was the left-branching complexity for that sentence. In other words, the left-branching complexity is the maximum number of simultaneously open left-branching dependencies (see Figure 1). Similarly, for right-branching complexity, at each word in each sentence, we calculated the number of dependents that would still connect to verbal heads that had been presented up to that word. The crucial difference between left and right-branching constructions is the order of heads and arguments. In left-branching dependencies the arguments precede the head. In right-branching sentences the heads are followed by the arguments further downstream in the sentence. The calculation of the right-branching complexity measure is thus symmetric with respect to the left branching complexity measure. Further details can be found in the supplementary material.

### 2.3 Task and Procedure

Within a measurement session, the stimuli were presented in a mini-block design, and alternated between a sentence block (5 sentences) and a word-list block (5 word-lists), for a total of 24 blocks. The type of starting block (sentences or word-lists) was counter-balanced across subjects. In order to check for compliance, 10 % of the trials were followed by a yes/no question about the content of the just presented sentence/word-list. Half of the questions on the sentences addressed the content of the sentence (e.g. *Did grandma give a cookie to the girl?)* whereas the other half, and all of the questions on the word lists, addressed one of the main content words (e.g. *Was the word ‘grandma’ mentioned?*). A substantial part of the questions on complex relative clause sentences concerned content from the relative clause. Based on the answers to the catch trial comprehension questions, subjects were defined as outliers (i.e., subjects with a negative distance > 1.5 times the interquartile range, from the mean) and were excluded from further analysis, as not compliant with the task.

For visual stimulus presentation, sentences/word-lists were presented word-by-word with a mean duration of 351 ms for each word (minimum of 300 ms and maximum of 1400 ms, depending on word length). Corresponding visual sentences and word-lists had the same total duration. The median duration of whole sentences/word-list was 8.3 s (range 6.2-12 s). Auditory sentences had a median duration of 4.2 s (range 2.8-6.0 s), and were spoken in a natural pace. The matching word-lists words were also read in a natural pace, with a brief pause between words, averaging 7.7 s (range 5.5-11.1 s) per word-list.

### 2.4 MRI data acquisition

The data were acquired with a SIEMENS Trio 3T scanner using a 32 channel head coil. We used a whole head T2*-weighted echo planar blood oxygenation level dependent (EPI-BOLD) sequence. The single-echo 2D ascending slice acquisition sequence (partial brain coverage) had the following specifications: Volume TR = 2.00 s, TE = 35 ms, 90 degree flip-angle, 29 oblique slices, slice-matrix size = 64 x 64, slice thickness = 3.0 mm, slice gap 0.5 mm, FOV = 224 mm, voxel size (3.5×3.5×3.0 mm^3^ during acquisition; interpolated to 2×2×2mm^3^ in SPM).

For the structural MR image volume, a high-resolution T1-weighted magnetization-prepared rapid gradient-echo pulse sequence was used (MP-RAGE; TE = 3.03 ms, 8 degree flip-angle, 1 slab, slice-matrix size = 256×256, slice thickness = 1 mm, field of view = 256 mm, isotropic voxel-size = 1.0×1.0×1.0 mm^3^).

### 2.5 Preprocessing

The primary imaging data were checked for quality, including checks of subject movement and signal drop out. The data were preprocessed with statistical parametric mapping software (SPM8; Welcome Trust Centre for Neuroimaging, London, UK, www.fil.ion.ucl.ac.uk/spm), which was also used for statistical analysis at the first-and second-level.

The first three EPI-BOLD volumes were removed to ensure T1-equilibrium. The remaining volumes were (1) realigned to correct for individual subject movement and (2) corrected for differences in slice-acquisition time.

The mean EPI-BOLD volumes were co-registered to the structural image (i.e., the structural image was the reference and the mean EPI-BOLD volume was the source images), and this transformation was then applied to all EPI-BOLD volumes. Structural images were spatially normalized to the structural image (T1) template provided by SPM8, using affine regularization. The transformation matrices generated by the normalization algorithm were applied to the corresponding functional EPI-BOLD volumes. All structural and functional images were spatially filtered with an isotropic 3D spatial Gaussian filter kernel (FWHM = 10 mm).

### 2.6 Statistical analysis

The fMRI data were analyzed statistically, using the general linear model framework and statistical parametric mapping (Friston et al. 2007) in a two-step mixed-effects summary-statistics procedure.

#### 2.6.1 First level models

The single-subject fixed effect analyses included a temporal high-pass filter (cycle cut-off at 128 s), to account for various low-frequency effects. All linear models included the six realignment parameters from the movement correction. In addition to two regressors, modeling the sentence and word-lists condition, all models included three regressors: the intertrial-interval, the inter-block-interval as well as the comprehension questions. This basic model was used to probe the language/sentence processing network (compared to a low-level baseline, more specifically the inter-block-interval) as an initial quality check. Each additional model as well as the contrasts are described in more detail below.

##### 2.6.1.1 Left vs. right-branching complexity

Regressors containing two sets (corresponding to sentences and word-lists, respectively) of five parametric sentence/word-list modulators, included: (1) number of letters; (2) number of syllables; (3) number of words; all of them referred to as ‘control regressors’; (4) the right-branching complexity; and (5) the left-branching complexity. These parametric modulators were entered as user-specified regressors, after convolution with the canonical HRF. To control for lexical level features, we associated each scrambled word-list version of a sentence with the left and right complexity measure of that sentence and contrasted complexity for sentences and word-lists. To verify the expectation that complexity should affect sentences and not unstructured word-lists, we also created contrasts probing the sentence and word-lists, separately, against the implicit baseline. We also formed the reverse contrasts (word-lists > sentences, i.e., negative effects of complexity) to those described in this paragraph.

##### 2.6.1.2 Left-branching processing complexity: temporal dynamics

In order to analyze activation changes over sentences, we divided each sentence/word-list presentation into four equally long time bins. For the time-bin analysis, parametric modulators were specified using pmod (parametric modulation) functionality implemented in SPM8. The order in which regressors are entered matters for this analysis, since each pmod regressor is orthogonalized with respect to the so called ‘unmodulated’ regressors as well as earlier pmod regressors (Mumford et al. 2015). Control regressors (i.e., numbers of words, letters and syllables), were thus entered before the regressors encoding the complexity measures, and the right-branching complexity measure was entered before the left-branching complexity measure, when assessing the effects of left-branching complexity; and conversely when assessing the effects of right-branching complexity.

We focused on the two ROIs that were most clearly and selectively responsive to left-branching complexity: the LIFG and LpMTG. To assess increases and decreases over the course of the sentences, we formed first-level linear contrasts across the complexity regressors corresponding to each time-bin (e.g., [−2 −1 1 2]).

Time courses were extracted from the LIFG and LpMTG ROIs, using the finite impulse response (FIR) model as implemented in MarsBar (Brett et al. 2002). The FIR model does not assume any particular shape of the hemodynamic response, but estimates the signal at each time bin (henceforth called time windows) separated by one TR (in our case 2 s), up to 24 s after stimulus onset, adjusting for effects of stimuli overlapping in time. Since our average visual stimulus duration was 8.3 s and our average ITI was 3.7 s, we analyzed the first seven time-windows, covering 0-14 s post-stimulus onset. We analyzed the differences in percent signal-change between high (left-branching complexity >=3) and low (left-branching complexity < 3) complexity, by introducing two sets of regressors (high/low) for sentences and word-lists in the FIR model. The differences in percent signal-change for high-low complexity were tested with a paired t-test at all time points. We corrected for multiple comparisons over seven time points using Bonferroni correction. In a follow-up analysis, we tested whether the complexity effects came in different time-windows for LpMTG, compared to LIFG. We compared the difference scores of the percent signal change for high complexity sentences – low complexity sentences, for LpMTG-LIFG, using one paired t-test.

#### 2.6.2 Second-level statistics and visualization

The second level comparison used a two-sample unequal-variance t-test. Visual and auditory data were modeled separately and then compared. Conjunctions were tested against the conjunction null (Nichols, 2005). Throughout the paper, statistics are reported with the following criterion for significance: presence of FWE-corrected voxels or clusters at *P*_FWE_ < .05, both in the whole brain analysis as well as in the ROI analysis.

In tables, all clusters significant at *P*_FWE_ < .05 on the whole brain level are reported including locations of voxels belonging to a cluster. Location and statistics of significant peak voxels (within or outside significant clusters) are also reported.

#### 2.6.3 ROI analysis

In the ROI-analysis, we used 10 mm spherical ROIs around meta-analytically established activation hotspots of semantic and syntactic unification, reported in Talairach space (Hagoort and Indefrey 2014). The two top coordinates for semantic and syntactic unification were close to each other, one pair in the LIFG and another in the LpMTG. We used the mean coordinate between the syntactic and semantic peak for our ROI-analysis. This resulted in the (Talairach space) coordinates [x y z] = [-45 20 10] for LIFG and [−50 −45 5] for LpMTG, which we then transformed to MNI space using Talairach Daemon (http://www.talairach.org/).

In addition, a LaSTG ROI was investigated in the left-vs right-branching complexity. This was again a 10 mm spherical ROI around the peak coordinate reported for the building of sentence structure in a related study (Brennan *et al.* 2012). These were [x y z] = [−50 11 −16] in MNI-space.

#### 2.6.4 Anatomical Inference

All local maxima are reported as MNI coordinates. Assignment of clusters and voxels to gyri, sulci and Brodmann’s areas (for LIFG) were done using the anatomy toolbox (version 21) implemented in SPM (Eickhoff et al. 2005).

## 3. RESULTS

### 3.1 Sentence processing vs. low-level baseline

We verified that our paradigm activated the classical perisylvian language network by comparing activation during sentence processing with a low-level baseline (i.e., fixation/rest inter-block-interval). A significant cluster spanning left anterior to posterior superior temporal cortices (middle and superior temporal gyrus), left inferior frontal gyrus (BA 44/45), the left inferior parietal lobe and left fusiform gyrus was observed (see Figure 2A; as well as Table S1 in the supplementary material). Two additional clusters (right superior temporal and right fusiform gyrus) as well as significant voxels in left orbitofrontal cortex, left precentral and left superior frontal gyrus, were also observed.

**Figure 2.**
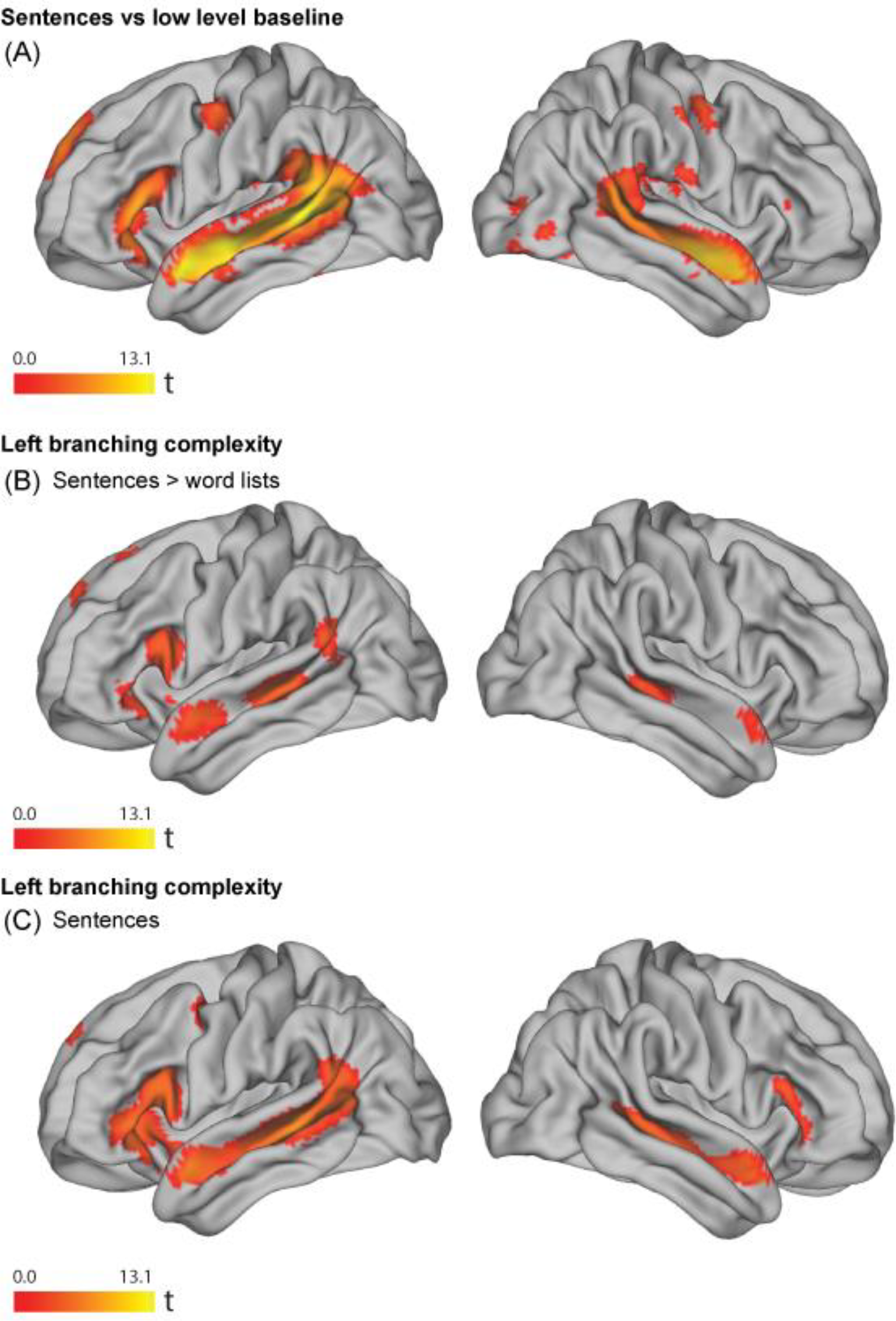
Results images were created based on the conjunction of data of the visual (N=102) and the auditory (N=102) presentation of the materials, using a threshold of *P* < .005 uncorrected, for the contrasts with significant results at the cluster or voxel level, using a criterion of *P*_FWE_ < .05. (A) Effect of sentences > low level baseline. (B) Effect of left-branching complexity in sentences > word-lists (word-list complexity is the complexity of the corresponding sentence). Peaks in a significant temporal cluster were observed in LpMTG and LaSTG. The LIFG ROI, indicated in red, contained a significant cluster. (C) Effect of left-branching complexity in sentences (i.e. without comparing with word-lists). Peaks in a large significant fronto-temporal cluster, were observed in LpMTG, LaSTG and LIFG. LpMTG: Left posterior middle temporal gyrus; LaSTG: Left anterior superior temporal gyrus; LIFG: Left inferior frontal gyrus.

### 3.2 Left-branching complexity

To investigate supramodal unification (Vosse and Kempen 2000; Hagoort 2005), we created conjunctions over visual and auditory contrasts of parametrically varied sentence complexity. We controlled for lexical features correlated with sentence complexity by subtracting corresponding word-lists parameters in the following way. Each scrambled word-list version of a sentence was associated with the left and right-branching complexity measure of that sentence. We analyzed the parametric effects of left-branching complexity for sentences compared to the corresponding effects for word-lists (henceforth referred to as sentences > word-lists contrasts).

There was a significant interaction between the parametric effect of left-branching complexity in sentences compared to word-lists throughout the left perisylvian language network (Figure 2B and Table 2). In this comparison, a significant cluster was observed in posterior middle/superior temporal cortices, anterior superior temporal gyrus, left inferior frontal gyrus (BA 44 and 45) and left inferior parietal cortex. There was no significant effects in the reverse comparison word-lists > sentences. To consolidate the findings in the sentence > word-list contrast, we verified that complexity had no effect on word-lists, by investigating the effect of left-branching processing load in sentences and word-lists separately. There were no significant effects of complexity in the word-list condition. In contrast, we observed a positive effect of left-branching complexity for sentences in the left perisylvian language network (Figure 2C and Table 2). There was no significant activity in the reverse comparison.

**Table 2.** Supramodal effect of left-branching complexity.

| Region | Cluster<br>size | Cluster<br>$P_{FWE}$ | MNI- coordinates | | | Voxel<br>$P_{FWE}$ | Voxel<br>$T_{201}$ |
| --- | --- | --- | --- | --- | --- | --- | --- |
|  |  |  | x | y | z |  |  |
| Left-branching * |  |  |  |  |  |  |  |
| Sentences > words |  |  |  |  |  |  |  |
| *LpMTG/LaSTG/LIFG/LIPL* | 2678 | .001 |  |  |  |  |  |
| *LpMTG/STS* |  |  | -52 | -30 | -4 | .003 | 5.13 |
| *LaSTG* |  |  | -56 | 2 | -16 | .042 | 4.47 |
| + *LIFG (BA 44,45), LIPL* |  |  |  |  |  |  |  |
| Left-branching |  |  |  |  |  |  |  |
| Sentences |  |  |  |  |  |  |  |
| *LpMTG/LaSTG/LIFG/LIPL* | 4225 | <.001 |  |  |  |  |  |
| *LpMTG/STS* |  |  | -50 | -36 | -2 | <.001 | 7.67 |
| *LaSTG* |  |  | -54 | 8 | -18 | <.001 | 6.40 |
| *LIFG (BA45)* |  |  | -48 | 28 | 0 | .001 | 5.19 |
| *LIFG (BA44)* |  |  | -56 | 18 | 14 | 0.003 | 5.15 |
| *LIPL I* |  |  | -50 | -54 | 16 | 0.011 | 4.82 |
| *LIPL II* |  |  | -54 | -56 | 18 | 0.013 | 4.77 |
| *LIPL III* |  |  | -52 | -54 | 12 | 0.019 | 4.68 |
| Right branching |  |  |  |  |  |  |  |
| Sentences |  |  |  |  |  |  |  |
| *ROI LaSTG 10mm, SVC* | 37 | .048 |  |  |  |  |  |
|  |  |  | -54 | 8 | -20 | .024 | 3.13 |
| No significant activations (neither at ROI-level) for: Left, Words > Sentences; Negative effect of Left, Sentences. |  |  |  |  |  |  |  |

Note: \*see supplementary methods for significant visual > auditory or auditory > visual activations.

### 3.3 Effects of left vs. right-branching complexity

For the corresponding comparisons for right-branching complexity to those reported in section 2.2, no significant effects were found for sentences > word-lists nor for word-lists > sentences. For the separate sentence contrast, there were no significant voxels or clusters. Our ROI analysis included three ROIs (LIFG, LpMTG and LaSTG). The LaSTG ROI contained a significant cluster for an increase in right-branching complexity. No significant effects were observed for word-lists nor for the corresponding comparisons in the reverse direction.

### 3.4 Left-branching complexity: temporal dynamics

Unification processes operate incrementally during word-by-word presentation of sentences. The unification load will thus vary over time, in particular for complex sentences. To describe the temporal dynamics of sentence unification, we divided each sentence and word-list into four time-bins of equal length. We focused on the left-branching complexity measure, since it showed the most prominent effects. For the ROI-analysis, we used the two ROIs that were most clearly active specifically to left-branching complexity: the LIFG and LpMTG. The right-branching complexity parameter and control parameters (number of words, number of syllables and number of letters) were entered into the model to ensure that we were studying the contribution of left-branching complexity to dynamic changes over time. We created linear contrasts probing monotonic increases and decreases in the parametric effect of the complexity measures across the four time-bins. We expected temporal dynamics related to sentence unification to be stronger in sentences compared to word-lists.

As shown in Figure 3A and Table 3, we observed a monotonic increase related to sentence unification in the LpMTG ROI. When analyzing sentences and word-lists separately, a monotonic increase over sentences was observed in both the LIFG and LpMTG ROIs. No decrease was observed.

**Table 3.** Supramodal left-branching complexity, *increase* across time-bins.

| Region | Cluste | Cluster | MNI- coordinates |  |  | Voxel | Voxel |
| --- | --- | --- | --- | --- | --- | --- | --- |
| | r size | $P_{FWE}$ | x | y | z | $P_{FWE}$ | $T_{201}$ |
| Left branching |  |  |  |  |  |  |  |
| Sentences > words |  |  |  |  |  |  |  |
| *ROI LpMTG 10mm, SVC* | 74 | .034 |  |  |  |  |  |
|  |  |  | -46 | -42 | 0 | .025 | 3.29 |
| Left branching |  |  |  |  |  |  |  |
| Sentences |  |  |  |  |  |  |  |
| *ROI LpMTG 10mm, SVC* | 242 | .029 | no significant peak voxels |  |  |  |  |
| *ROI LIFG 10mm, SVC* |  |  | -50 | 20 | 16 | .028 | 3.25 |
| No significant activations (neither at ROI-level) for: Any of the corresponding decreases; Right, Sentences > words; Right, Sentences. Furthermore, there were no effects of modality (visual > auditory or auditory > visual). |  |  |  |  |  |  |  |

**Figure 3.**
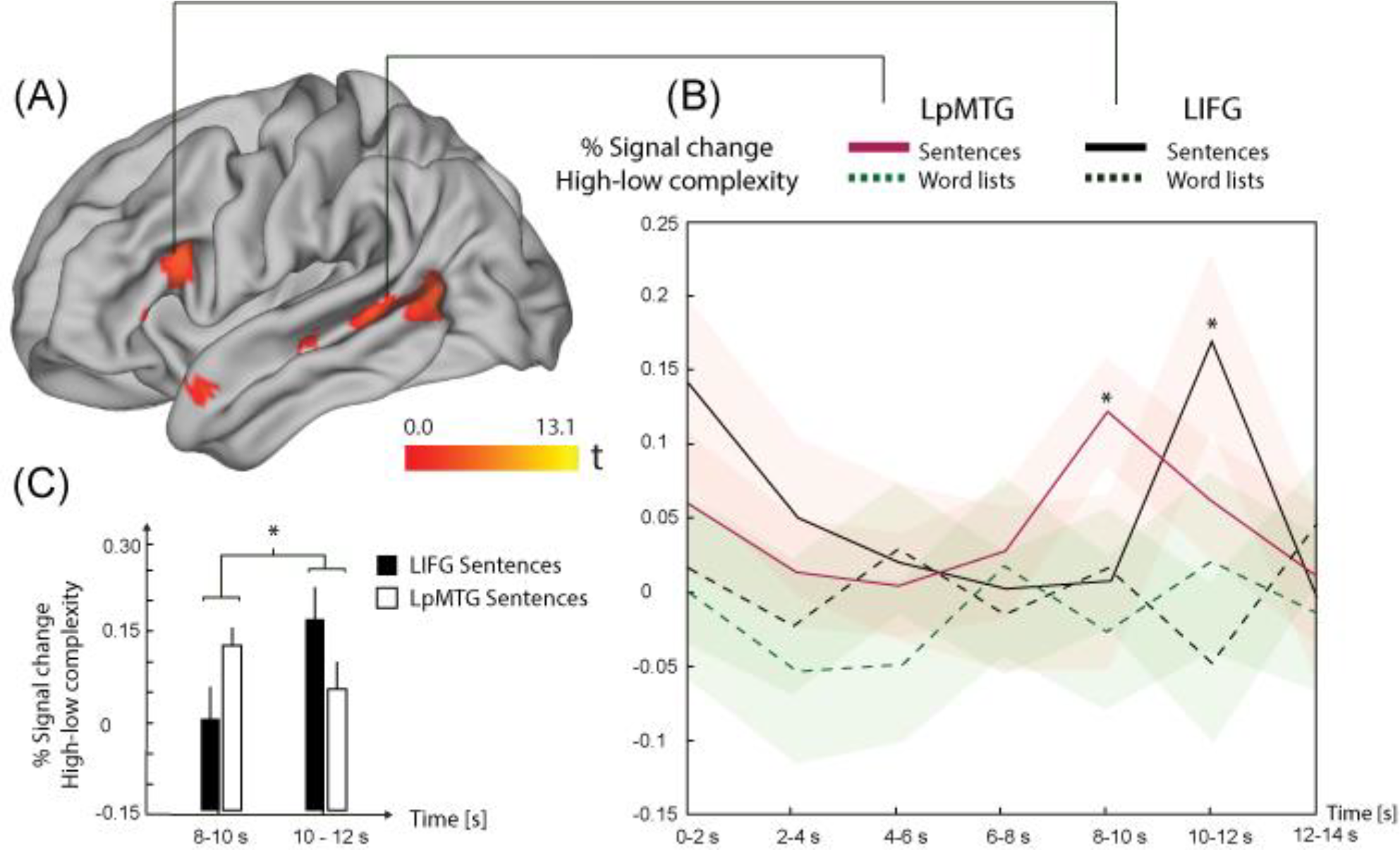
(A) In red, left-branching complexity, increase across time-bins for sentences. Results images were created based on the conjunction of the data for visual and auditory presentations of the materials, using a threshold of *P* < .005 uncorrected, for the contrasts with significant results at the cluster or voxel level, using a criterion of *P*_FWE_ < .05. In the LpMTG ROI, there was a significantly (using SVC) steeper increase for sentences > word-lists. (B) To illustrate the observed dynamics in (A) further, with time courses, we focused on the written sentences and word lists, which has longer presentation times for sentences (compared to the naturally spoken sentences) and thus provides more robust estimates of changes in the BOLD response over the sentence. Complexity difference scores (high-low) of % signal change from LIFG and LpMTG ROIs, in sentences and word-lists are plotted over seven time-windows. Red shaded areas represent +/− 1 SEM around each data point for sentences and likewise for word-lists, in green shade. The % signal change is relative to the mean activity of this ROI in the whole session, thus, note that the first time point (0-2 s) can be active since already a few words were presented during this time. The complexity effect within sentences was significant at 8-10s s in LpMTG (T_101_ = 3.28, *P*_Bonf_ = .01) and at 10-12 s in LIFG (T_101_ = 2.82, *P*_Bonf_ = .04). Both of these survive Bonferroni correction for testing seven time points. The seventh time-point (12-14s) contains the inter trial interval, on average. All four mean time courses returned to the session mean in at this time-point. (C) The complexity effect for sentences occurred in an earlier time-window for LpMTG, compared to LIFG. There was a significant interaction between the sentence complexity effect in LpMTG vs LIFG, at 8-10 s vs 10-12 s (T_101_ = 2.39, *P* = .02).

To follow-up on this finding, we extracted percent signal change time courses (using a finite impulse response model) from the LIFG and LpMTG ROIs, in the visual group (Figure 3B). The visual sample has longer presentation times compared to the naturally spoken sentences. Thus, more robust estimates of changes in the BOLD response over the sentence are expected for this part of the sample. We analyzed the differences in percent (%) signal change between high (left-branching complexity >=3, 168 sentences) and low (left-branching complexity < 3, 192 sentences) complexity, as a function of time. The complexity effect (high-low) for sentences was significant at 8-10 s in the LpMTG ROI (T_101_ = 3.28, *P*_Bonf_ = .01) and at 10-12 s in the LIFG ROI (T_101_ = 2.82, *P*_Bonf_ = .04).

We Bonferroni corrected for testing seven time points. The complexity effect for sentences thus occurred in an earlier time-window in LpMTG than in LIFG. There was a significant difference between the sentence complexity effect in LpMTG - LIFG, at 8-10 s vs 10-12 s (T_101_ = 2.39, *P* = .02, see Figure 3C). The seventh time-point (12-14s) contains the inter-trial-interval.

## 4. DISCUSSION

Our study provides support for the hypothesis that unification should be understood as a supramodal process, common to sentences presented as speech or text. The current results show that the supramodal network subserving unification includes the LIFG, the bilateral MTG (both anterior and posterior parts) as well as LIPL. We observed an effect of left-branching complexity in the perisylvian language regions, irrespective of modality. Moreover, we observed an absence of modality specific effects related to unification processes (i.e. no effects when comparing visual > auditory or auditory > visual samples). This finding provides further support for unification as a process that is not modality-specific. Our results thus support a view where the visual and auditory input streams converge in a supramodal unification network, most clearly in LIFG and LpMTG.

Previous studies, comparing complex vs simple sentences, have implicated the inferior frontal and temporal heteromodal association cortices as supramodal regions associated with sentence structure manipulations (Michael *et al.* 2001; Constable *et al.* 2004; Shankweiler *et al.* 2008; *Braze et al.* 2011). The large sample-size and the use of parametrical variation of processing complexity increase both sensitivity and validity compared to these previous studies. Our results support the view that LIFG and the bilateral MTG (both anterior and posterior parts) are key nodes of the supramodal unification network. Moreover, our results suggest that additional regions (e.g. LIPL, see further discussion below) could be considered as a part of this network (Binder et al. 2009).

The effects of left-branching complexity show increased activation levels in mainly left perisylvian language regions for syntactically more complex sentences. Moreover, the results show that the effect of left-branching processing complexity in LIFG and LpMTG increased over the course of the sentence. The corresponding changes were non-significant for LIPL, and similar sentence progression effects were not observed for the right-branching structures. Thus, in summary, left-branching complexity effects are stronger and show different dynamics over the sentence compared to the right-branching complexity effects. This highlights the fact that the neuronal underpinnings of sentence complexity cannot be fully appreciated without understanding which sentence processing operations a complexity measure taps into. Another good example of this approach can be found a recent ECoG study on visual sentence processing (Nelson et al. 2017; see also Fedorenko et al., 2016). By using many processing models to predict neural activity over the sentence, they found evidence for a structure building process in LIFG and posterior superior temporal cortices. It is possible that a substantial part of the variance in the neural dynamics explained by our left-branching complexity measure is shared with their word-by-word increases in high gamma for every word in a constituent. Our results complement their results in six crucial ways: (1) by showing a functional asymmetry in neural processing of dependencies that go from the head to the dependent (i.e. left-branching) compared to the other way around; (2) by showing the spatial specificity of these effects with FMRI; (3) by a 15 fold increase in the sample size to ensure robust effects; (4) by increasing the variability of the sentence structures used, since Nelson et al. 2017 restricted their sentence materials to only a single sentence structure; (5), by using a more realistic dependency grammar rather than binary branching trees; and finally (6) by extending these results from the visual modality only to the auditory modality as well, showing that the effects of left-branching complexity are likely to be supramodal. All modality comparisons (more extensively reported in the supplemental results) support this conclusion. Below we interpret the results by focusing on the operations targeted by our left-branching complexity measure, using the effects of right-branching complexity for contrast and comparison.

A crucial aspect of language processing is the binding of the head of a verb phrase to its arguments. At least two factors determine processing complexity of unifying a verb phrase. One is the position-distance between the head and its arguments in a sentence. The second is the order of heads and arguments. There are several possible reasons for this, such as an additional processing cost of storing unbounded arguments, as well as an asymmetry in predictive power (the head is a stronger prior for arguments than vice versa). In our study we computed a complexity measure for each individual sentence in the experiment, based on a formal characterization of the processing consequences of simultaneous unbounded arguments. This complexity measure modulated activity in LIFG and LpMTG in a parametric way, strongly suggesting that these regions together play a crucial role in the unification of lexical elements (i.e., words) into an overall structural sentence configuration.

When studying the activation dynamics across the sentence in the different nodes of the supramodal unification network, we observed a particularly interesting pattern of results in LIFG and LpMTG. In these regions, activity increased over the sentence for left-branching structures. No such effect was observed in LIPL (see Figure 3). We calculated the maximum point of simultaneous unification of unbounded arguments (usually referred to as dependents for other phrases (e.g., NP) than VPs), which occurred on average around half-way into our sentences. Furthermore, the left-branching complexity measure indexes sentences with many simultaneous non-attached constituents, a cost which increases with each non-attached constituent presented, until a culmination point around half-way into the sentences. Since the priors for unification are relatively low in left-branching dependencies compared to right-branching dependencies, increased activation is expected towards the later parts of the sentence, given the increased number of unbounded arguments.

An additional novel finding is that the complexity effect had an earlier time course in LpMTG than in LIFG. Both areas contribute to syntactic processing (Hagoort & Indefrey, 2014), but the effect in temporal cortex might be more local, related to the verbal head and argument binding, whereas the complexity effect in LIFG might be more global, presumably related to the consequences for computing the overall phrasal configuration of the whole sentence.

Right-branching complexity produced stronger activation than left-branching complexity in LaMTG. This could be due to the stronger expectations about upcoming lexical-semantic information in the former case. This view on LaMTG contribution to structure building fits well with the current views on the anterior temporal lobe function within linguistic and conceptual processing more generally (Patterson et al. 2007). There is furthermore evidence that conceptual combinations are processed in the anterior temporal lobe (e.g. Baron & Osherson, 2011). In line with our view, Pallier, (2011) only observed structure building (as indexed by an increasing response to increasing constituent size) in LaMTG for materials with real words and not in a Jabberwocky condition, where only posterior temporal and inferior frontal regions were observed (see also Goucha and Friederici 2015).

The measure of simultaneous non-local dependencies that we used correlates with the presence of non-local dependencies as well as their total length (see supplementary material). Our results thus show the neural underpinnings of the ubiquitous difficulty observed with processing non-adjacent dependencies (Gibson 1998; Grodner and Gibson 2005). In addition, our results reflect the increased difficulty of unifying complex left-compared to right-branching sentence aspects, at least in the context of a language like Dutch, which features considerable proportions of both. Our results suggests that *simultaneity* (or overlap) of multiple unresolved non-adjacent dependencies, rather than linear distance of non-adjacent dependencies, is a major factor contributing to the difficulty of processing non-adjacent dependencies (see additional analyses supporting this conclusion in section S2.2 in the supplementary material). This highlights the importance of understanding which sentence processing operation(s) a complexity measure taps into.

In conclusion, we have characterized the basic spatiotemporal dynamics of a supramodal unification network consisting of the LIFG, bilateral MTG and the LIPL. The finding that activity in this network increases over the sentence in this network is novel (see possibly related oscillatory modulations in ECoG and MEG, respectively: Fedorenko et al. 2016; Lam et al. 2016). These results support our interpretation that binding of unbounded arguments increase towards the end of the sentence, resulting in stronger activations in LIFG and LpMTG. We also show the neural underpinnings of the ubiquitous difficulty observed with processing overlapping non-adjacent dependencies is independent of modality. Both listening to speech and reading activate the same neuronal circuitry once the modality-specific input is mapped onto lexical items, in the service of the brain’s capacity to make sense beyond the processing of single words.

## ACKNOWLEDGEMENT

We thank Laura Arendsen, Manuela Schuetze, Tineke de Haan, and Charlotte Poulisse for assisting with stimuli construction, participant recruitment, data collection and preprocessing.

## Author contributions

J.U., A.H., P.H. designed research. J.U., A.H., N.L. performed research. J.U., J-M.S, K.H., G.K., K.M.P., A.vdB. contributed to analytic tools. J.U. analyzed data. J.U., P.H. wrote the paper.

