## Supplement for "Supramodal Sentence Processing in the Human Brain: Fmri Evidence for the Influence of Syntactic Complexity in More Than 200 Participants"

### S1. Supplementary methods

#### S1.1 Participants

All participants were right-handed as assessed by the Bever handedness questionnaire (including familial handedness), had normal or corrected-to-normal vision, and reported no history of neurological, developmental or language deficits. We screened for medication use and excluded anyone with medication on prescription. The instructions further contained a statement that no medication, alcohol or drugs should be used on the day of the measurement. 38 participant were excluded because of technical problems (20) (the most common were problems with data transfer, scanner hardware errors, presentation software problems resulting in faulty triggers or absence of comprehension questions), poor data quality due to excessive blinking (in the MEG session, affecting the amount of remaining MEG trials after artefact rejection) or more than 3.5 mm movement (4), study interruption (6), not fulfilling the inclusion criteria (full list: 18-35 years, right-handed, self-reported Dutch monolingual language background, normal or corrected-to-normal vision, no self-reported history of neurological, developmental or language deficits, MRI-compatibility (not pregnant, no claustrophobia, no incompatible devices, no incompatible tattoos, no incompatible self-reported operation history or metal in body) (4), or since participants had poor compliance with the task (4) (removal of these four outliers in task performance meant that all included participants had more than 59% correct, mean 80% correct +/- standard deviation of 9%). A total of 204 participants remained after exclusion of these 38 participants.

#### S1.2 Recording and post-processing of auditory stimuli

The audio files were recorded in stereo at 44100 Hz. During the post processing the audio files were low-pass filtered at 8500 Hz and normalized so that all audio files had the same peak amplitude, and same peak intensity. In the word-list condition, each word was separated by 300 ms of total silence. The transition from silence to speech was ramped at the onset (rise-time of 10 ms) and offset (fall-time of 10 ms) of single words in the word-list condition, and for sentence onset.

#### S1.3 Practice task

Prior to the task, participants read a written instruction of the task and asked questions for clarification. Furthermore, the experimenter emphasized that the sentences and word-lists should be attended carefully, and discouraged attempts to integrate the words in the word-list condition. Finally, to familiarize the participants with the task, they did a practice task with stimuli separate from the actual experiment.

#### S1.4 Stimulus presentation and comprehension questions

Each participant was presented with 180 sentences and 180 word-lists, either entirely in the visual or auditory modality. These stimuli where divided into three subsets, such that each participant saw 2/3 of the stimuli set in the MEG (120 trials of each condition) and 1/3 in the fMRI (60 trials). Across participants, each subset was presented as many times in MEG as in fMRI.

Each participant was presented with each stimulus once, in either the sentence or the word-list condition, but not in both. The presentation of the sentence and word-list versions of the items was counterbalanced across subjects. The auditory versions of the stimuli were recorded by a female native Dutch speaker. The word-lists were pronounced with a neutral prosody and a clear pause (300ms) between each word.

To indicate sentence- and word-lists blocks, the start of each block began with a 1500 ms block type indication (*zinnen* (sentences) or *woorden* (words)). Only in sentences did the first word begin with a capital letter, while the last word ended with a full stop. The inter-trial interval was jittered between 3200 – 4200 ms. During this period, an empty screen was presented, followed by a fixation cross.

The visual stimuli were presented with a LCD projector (vertical refresh rate of 60 Hz) situated outside the fMRI scanning room, and projected via mirrors onto the screen inside the measurement room. All stimuli were presented in a black mono-spaced font on a gray background at the center of the screen within a visual angle of 4 degrees using the Presentation software (Version 16.0, Neurobehavioral Systems, Inc.). For auditory stimulus presentation, sounds were presented via plastic tubes and ear pieces to both ears. Before the experiment, the hearing threshold was determined individually, and the stimuli were then presented at an intensity of 50 dB above the hearing threshold, with the obtained volume individually pretested on top of the EPI-sequence noise, to verify that all stimuli were clearly audible.

Each word was separated by an empty screen for 300 ms before the onset of the next word. The presentation time of each word was weighted by the number of letters in the word, providing a natural reading experience. In this way reading of the words was also matched to some extent to the total duration of the audio-version of the stimuli. For any given sentence (or word-list) the variable presentation duration of a single word was a function of the following quantities: (i) the total duration of the audio-version of the sentence/word-list (audiodur), (ii) the number of words in the sentence (nwords), (iii) the number of letters per word (nletters), and (iv) the total number of letters in the sentence (sumnletters). Specifically, the duration (in ms) of a single word was defined as: (nletters/sumnletters)*(audiodur+2000-150*nwords). The minimum duration of short words was set to 300 ms irrespective of the relative weighting described by the formula. Note that this formula does not apply to the auditory sentences, which were spoken in a natural pace.

Half of the comprehension questions addressed the content of the sentence (e.g. *Did grandma give a cookie to the girl?),* whereas the other half, and all of the questions on the word-lists, asked to verify the presence of a content word (e.g. *Was the word 'grandma' mentioned?*). Half of the questions on complex relative clause sentences concerned the content from the relative clause, to make sure the participants comprehended all parts of the sentence. Subjects answered the question by pressing a button for 'Yes'/ 'No' with their left index and middle finger, respectively. The outlier procedure described in the main methods section resulted in a threshold of 59% percent correct, which all included subjects passed.

**S1.5 Dependency trees: heads, dependents and the complexity measure**

An important aspect of the syntactic complexity of the sentences can be formalized in terms of their dependency structure. The automatic FROG-parser (http://antalvandenbosch.ruhosting.nl/) was used to create a dependency tree for every sentence (see section S1.5). The resulting trees were manually checked and corrected.

Like the more familiar phrase-structure trees, dependency trees encode syntactic aspects of word groups (phrases), but the aspects they emphasize differ. Phrase-structure trees specify the subphrases that a phrase is composed of, for instance, a Verb Phrase consisting of a Verb followed by a Noun Phrase. Phrases typically include one word, called the Head, that is more important than the other subphrases, called Dependents. In the example, the Verb functions as Head because it imposes properties on the Noun Phrase rather than vice-versa: It assigns accusative case to the Noun Phrase. Phrase-structure and dependency trees are similar because properties encoded by the former usually can be derived from the latter, and vice-versa. We have opted for dependency trees for a practical reason—the availability of a high-quality dependency parser for Dutch.

In Dutch, the finite verb is considered to be the head, in the main clause as well as in all types of subordinate clauses (complement, adverbial, relative). In clauses with an auxiliary, this auxiliary is the head of the clause, not the non-finite verb depending on the auxiliary. These definitions may not all correspond to English. There, a subordinating conjunction is often considered to be the head of an adverbial or complement clause. Sometimes, the auxiliary (as suggested by the name) is viewed as depending on the full verb, because the latter verb carries most of the meaning. This is however a semantic, not a syntactic criterion.

In order to calculate the right branching sentence complexity measure, we took the maximum of the word by word right branching complexity, as the right-branching complexity of that sentence. Thus, in other words, the right-branching complexity measure is the maximum number of simultaneously open right-branching dependencies.

For both left and right-branching complexity calculations, we focused on verbal heads since they are heading clauses. Clausal structure is more encompassing than the more local structure of phrases headed by other parts of speech (e.g., noun phrases, prepositional phrases, adjectival phrases). To verify this on our own material, we compared the left-branching complexity between sentences with and without relative clauses, for verbal and non-verbal heads. For verbal heads, left-branching complexity was higher for sentences with relative clauses (mean = 3.0 +/- 0.7 standard deviations) than sentences without (mean = 1.4 +/- 0.9 standard deviations). For non-verbal heads, there was no significant difference between sentences with relative clauses (mean = 1.9 +/- 0.5 standard deviations) and those without (mean = 2.1 +/- 0.5 standard deviations). It is assumed that clausal structure exerts the main influence on cognitive complexity. Note that although our complexity measures are designed to index *processing* complexity, for the sake of brevity we will refer to the outcome of calculations using these measures as “left/right-branching complexity”.

### S1.6 Preprocessing

Co-registration of the structural and functional images was checked for each individual subject by displaying the structural and the first functional image with SPM’s checkreg. Normalization was checked by displaying the structural image, the functional image and the template, again using checkreg. No deviant co-registration or normalization was found.

### S1.7 Left vs right-branching complexity

In order to compare the effects of left- and right-branching complexity directly, we analyzed a subset of the stimulus materials, selecting 320 of the 360 sentences, with their word-lists versions. We removed the sentences that had a combination of a high left-branching and a low right-branching complexity. In the resulting subset, the left- and right-branching complexity did not differ significantly over sentences (Wilcoxon test; ranksum: 103984, Z = .64, $P$ = .52, thus balancing the stimulus set for the left- vs. right-branching comparisons). We performed comparisons of left- vs. right-branching complexity on this stimulus set, by creating the same model as described in the main methods section.

Similarly, in order to compare the dynamic effects for left and right-branching processing load (directly), we used the same methods as in section (2.6.1.1). This model was implemented in SPM12, without orthogonalization between pmods. Thus, the order of entering pmods does not matter.

#### S1.8 First level model: total dependency length

An identical model to that used to investigate the left and right-branching complexity was used, but instead of these measures, we now used the total dependency length measure.

### S2. Supplementary results

#### S2.1 Left vs right-branching complexity

We analyzed the difference between the left and right-branching complexity directly. To do so, we selected a subset of the stimulus material, 320 of the 360 sentences, and their word-lists versions. In this subset, the left and right-branching complexity measures did not differ over sentences, as measured with a Wilcoxon rank test (rank sum: 103984, Z = .64, $P$ = .52). When contrasting left > right-branching complexity, for sentences > word-lists, a significant effect was observed in the LIFG ROI. In addition, analyzing the sentences and word-lists separately, there was a left-branching > right-branching complexity effect for sentences in the LpMTG ROI. No corresponding effects where observed for right-branching > left-branching complexity, and there were no effects of left-branching > right-branching or right-branching > left-branching complexity in word-lists (see Table S1). Similarly, we tested the left-branching vs right-branching complexity effects in the dynamic changes across timebins (see Table S2). There was an effect for left-branching > right-branching complexity, for the increase across timebins, in the LpMTG locus, both when looking at sentences separately and when contrasting sentences to word lists.

**Table S1**

Supramodal left vs right-branching complexity.

| Region | Cluster size | Cluster $P_{FWE}$ | MNI- coordinates x | y | z | Voxel $P_{FWE}$ | Voxel $T_{201}$ |
|---|---|---|---|---|---|---|---|
| Left > Right branching* Sentences > words | | | | | | | |
| *ROI LIFG 10mm, SVC* | | | -40 | 20 | 4 | .045 | 2.88 |
| Left > Right branching*,** Sentences | | | | | | | |
| *ROI LpMTG 10mm, SVC* | 59 | .042 | | | | | |
| | | | -52 | -38 | 0 | .036 | 3.00 |

No significant activations at whole brain or ROI-level for: Right, Sentences > words*; Right > Left*, Sentences > words; Right > Left*, Sentences; Right, Words > Sentences; Negative effect of Right, Sentences.

Note: * The direct comparisons of left vs right-branching complexity were performed on a subset of 320 (out of 360) sentences. In this subset, the left and right-branching complexity measures did not differ significantly over sentences ($p > .50$). SVC: small volume correction

**see supplementary methods for significant visual > auditory or auditory > visual activations.

**Table S2**

Supramodal left- vs right-branching complexity, *increase* across time-bins.

| Region | Cluster size | Cluster $P_{FWE}$ | MNI- coordinates x | y | z | Voxel $P_{FWE}$ | Voxel $T_{201}$ |
|---|---|---|---|---|---|---|---|
| Left > Right branching* Sentences > words | | | | | | | |
| *ROI LpMTG 10mm, SVC* | 83 | .027 | | | | | |
| | | | -46 | -42 | 2 | .011 | 3.63 |
| Left > Right branching* Sentences | 92 | .022 | | | | | |
| | | | -46 | -42 | 2 | .010 | 3.70 |

No significant activations (neither at ROI-level) for: Any of the corresponding decreases; Right > Left*, Sentences > words, Right > Left*, Sentences. Furthermore, there were no effects of modality (visual > auditory or auditory > visual).

Note: *The direct comparisons of left vs right-branching complexity were performed on a subset of 320 (out of 360) sentences, turning off orthogonalization between pmods in SPM12. In this subset, the left and right-branching complexity measures did not differ significantly over sentences ($p$ > .50). SVC: small volume correction

**Table S3**

Supramodal effect of Sentences vs low level baseline (fixation/rest).

| Region | Cluster size | Cluster $p_{FWE}$ | MNI- coordinates x | y | z | Voxel $p_{FWE}$ | Voxel $T_{201}$ |
|---|---|---|---|---|---|---|---|
| Sentences > IBI | | | | | | | |
| *LpMTG/LaSTG/LIFG/LIPL/ Left Fusiform G* | 7265 | <.001 | | | | | |
| *LMTG (mid)* | | | -56 | -10 | -12 | <.001 | 16.21 |
| *LpMTG/STS* | | | -54 | -46 | 10 | <.001 | 15.75 |
| *LaSTG* | | | -48 | 14 | -22 | <.001 | 14.64 |
| *Left Fusiform G I* | | | -36 | -38 | -18 | <.001 | 7.85 |
| *Left Fusiform G II* | | | -30 | -34 | -16 | <.001 | 7.67 |
| *Thalamus* | | | -8 | -28 | -2 | <.001 | 7.53 |
| *LIFG (BA 45)* | | | -56 | 28 | 8 | <.001 | 7.52 |
| *LIFG (BA 47)* | | | -40 | 32 | -10 | <.001 | 6.94 |
| *L Medial TL I* | | | -40 | -16 | -20 | 0.002 | 5.33 |
| *L Medial TL II* | | | -22 | -14 | -14 | 0.014 | 4.86 |
| + *LIPL* | | | | | | | |
| *RpMTG/RaSTG* | 2909 | <.001 | | | | | |
| *RaSTG* | | | 48 | 14 | -20 | <.001 | 11.92 |
| *RMTG (mid)* | | | 54 | -6 | -14 | <.001 | 11.73 |
| *RpMTG/STS* | | | 50 | -36 | 6 | <.001 | 7.66 |
| *R Rolandic Operculum* | | | 40 | -20 | 20 | 0.037 | 4.61 |
| *R Fusiform G* | 1075 | .012 | | | | | |
| *R Fusiform G posterior* | | | 30 | -52 | -10 | .001 | 5.50 |
| *R Fusiform G anterior* | | | 28 | -30 | -18 | .001 | 5.49 |
| *Left orbitofrontal gyrus* | | | -6 | 36 | -18 | <.001 | 7.15 |
| *LSFG* | | | -10 | 58 | 32 | <.001 | 6.29 |
| *Left precentral gyrus* | | | -42 | -6 | 48 | <.001 | 5.76 |

**S2.2 Modality specific effects: Left vs. right-branching complexity**

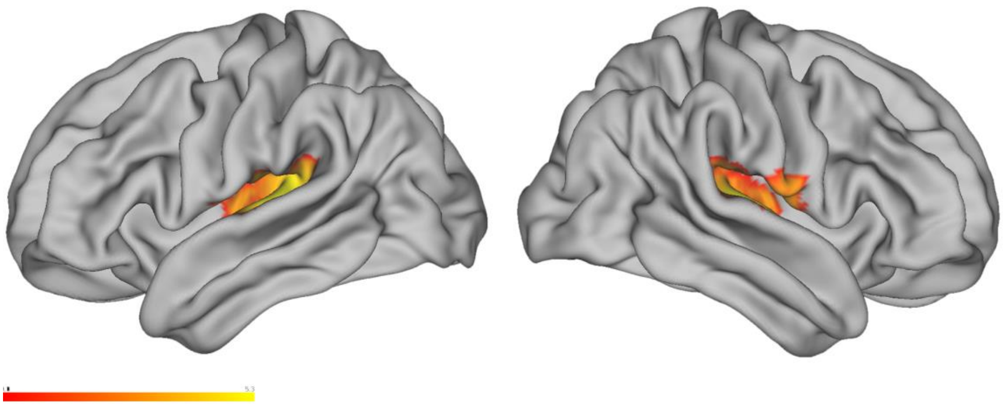

Figure S1. Left-branching, Sentences.Visual > Auditory (*P* < .005 uncorrected).

Here we report all modality specific effects (visual > auditory) and (auditory > visual) effects that were present in some of the contrasts reported in results sections 3.2 and forward. There were visual > auditory effects for the left-branching complexity measure, for sentences, and when comparing left > right-branching complexity, for sentences > word-lists, again in the visual > auditory direction (Figure S1 and S2, Table S4). In a follow up analysis, we tested whether the observed effects in these visual > auditory contrasts were due to an effect of left-branching complexity in the opposite/negative direction (i.e. higher BOLD response for lower complexity) for the auditory group. This was the case (Table S4).

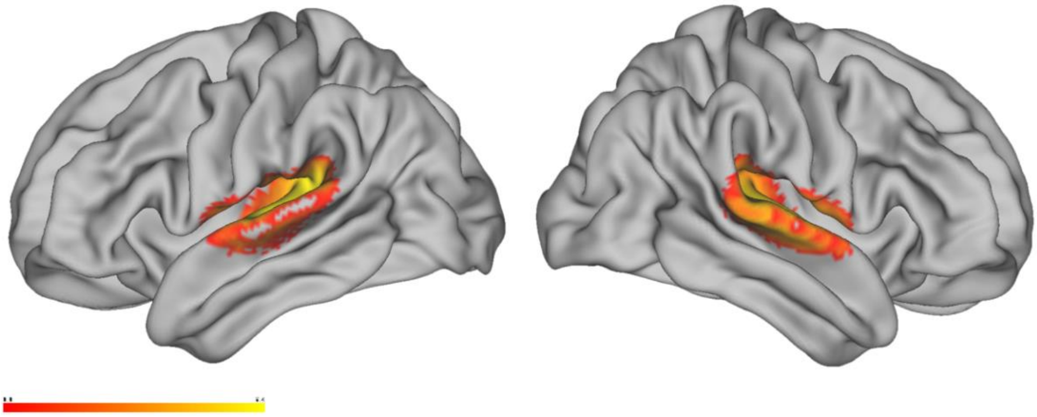

Figure S2. Left > Right, Sentences > Word-lists. Visual > Auditory (*P* < .005 uncorrected).

**Table S4**

Modality specific effects left (vs right) complexity in sentences vs word-lists

| Region | Cluster size | Cluster $P_{FWE}$ | MNI- coordinates x | y | z | Voxel $P_{FWE}$ | Voxel $T_{201}$ |
|---|---|---|---|---|---|---|---|
| Left-branching Sentences Visual > Auditory | | | | | | | |
| *LSTG/Heschl's gyrus* | 1575 | .011 | | | | | |
| | | | 48 | -16 | 8 | <.001 | 5.74 |
| | | | 44 | -24 | 10 | .001 | 5.44 |
| *RSTG/Heschl's gyrus* | 1764 | .009 | | | | | |
| | | | -38 | 30 | 12 | <.001 | 5.73 |
| | | | -44 | -24 | 12 | <.001 | 5.59 |
| *Left Subiculum* | 1221 | .031 | | | | | |
| Right-branching, Sentences > word-lists Auditory > Visual | | | | | | | |
| *ROI LIFG 10mm, SVC* | 36 | .048 | -52 | 20 | 10 | .003 | 2.96 |
| Left vs right-branching Sentences > word-lists Visual > Auditory | | | | | | | |
| *LSTG* | 2954 | <.001 | | | | | |
| | | | -50 | -18 | 4 | <.001 | 7.32 |
| | | | -40 | -30 | 10 | <.001 | 6.38 |
| *RSTG* | 2429 | .001 | | | | | |
| | | | 54 | -18 | 6 | <.001 | 6.57 |
| | | | 56 | -8 | 2 | <.001 | 6.24 |

No significant activations for visual > auditory or auditory > visual in: Left, sentences > word-lists; Right, sentences; Left vs right, sentences. No significant activations for auditory > visual in: Left, Sentences; Left vs right, sentences > word-lists. No significant activations for visual > auditory in: Right, sentences > word-lists.

**Table S5**

Modality specific auditory effect of left (vs right) branching complexity.

| Region | Cluster | Cluster | MNI- coordinates | | | Voxel | Voxel |
|---|---|---|---|---|---|---|---|
| | size | $P_{FWE}$ | x | y | z | $P_{FWE}$ | $T_{201}$ |
| Left branching<br>Negative direction<br>Sentences<br>Auditory | | | | | | | |
| *LSTG* | 1988 | .004 | | | | | |
| | | | -38 | -30 | 12 | <.001 | 6.16 |
| | | | -48 | -20 | 8 | <.001 | 5.81 |
| *RSTG* | 1741 | .007 | | | | | |
| | | | 50 | -16 | 8 | <.001 | 6.10 |
| *Left fusiform gyrus* | | | -32 | -38 | -10 | .009 | 4.87 |
| Left > right-branching<br>Negative direction<br>Sentences > word-lists<br>Auditory | | | | | | | |
| *LSTG/Heschl's gyrus* | 5217 | <.001 | | | | | |
| | | | -50 | -20 | 6 | <.001 | 9.18 |
| | | | -40 | -30 | 10 | <.001 | 7.74 |
| *RSTG* | 3131 | <.001 | | | | | |
| | | | -54 | -18 | 6 | <.001 | 8.05 |

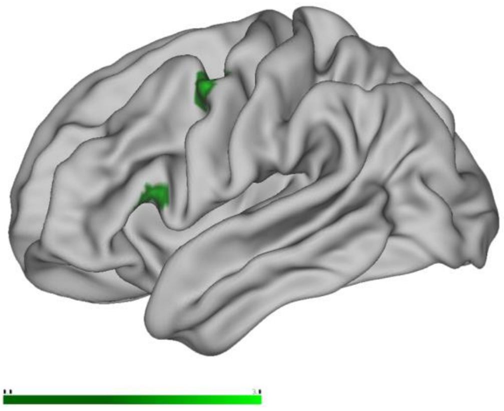

Figure S3. Right-branching, Sentences > Word-lists. Auditory > Visual (*P* < .005 uncorrected).

### S2.2 The total dependency length complexity measure

The left-branching complexity measure is each sentence's maximum number of simultaneously open left-branching dependencies. For comparison of the complexity measures we use with the total dependency length measure (Futrell et al. 2015). A universal tendency for minimizing total dependency length was recently reported (Futrell *et al.* 2015). We calculated the total dependency length for our sentences. Using partial correlations controlling for number of letters, words and syllables, the total dependency length measure was robustly correlated with our left-branching complexity measure ($\rho_{(rho)}$ = .58, $P < .001$), but not with right-branching complexity measure ($\rho_{(rho)}$ = -.03, $P = .52$). We analyzed the total dependency length in a separate model, assessing effects of neural infrastructure subserving processes of maintenance of lexical items, as a complement to the analysis of left-branching complexity. We observed no significant effect of total dependency lengths in the sentences > word-lists comparison. However, we observed an effect of increasing total dependency length in the LpMTG ROI, for the separate sentence contrast (Figure S4 and Table S6). There were modality specific effects, in the direction of auditory > visual (see Figure S5 and Table S7). We followed up by analyzing the visual and auditory groups separately, the auditory sample had significant clusters for total dependency length, in both sentences > word-lists and sentences when analyzed separately (Figure S5 and Table S7).

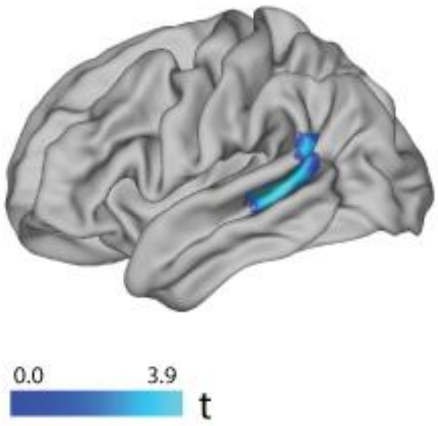


Figure S4. Positive parametric effect of total dependency length in sentences, $P < .005$ uncorrected. There were significant clusters and voxels in the LpMTG ROI. Conjuction over auditory and visual groups.

**Table S6**

Supramodal effect of total dependency length (TDL) in sentences and word-lists.

| Region | Cluster size | Cluster $p_{FWE}$ | MNI- coordinates x | y | z | Voxel $p_{FWE}$ | Voxel $T_{201}$ |
|---|---|---|---|---|---|---|---|
| TDL, Sentences | | | | | | | |
| *ROI LpMTG 10mm, SVC* | 279 | .011 | | | | | |
| *ROI LpMTG 10mm, SVC* | | | -50 | -40 | 2 | .001 | 4.13 |
| No significant activations (neither at ROI-level) for TDL, Sentences > words | | | | | | | |

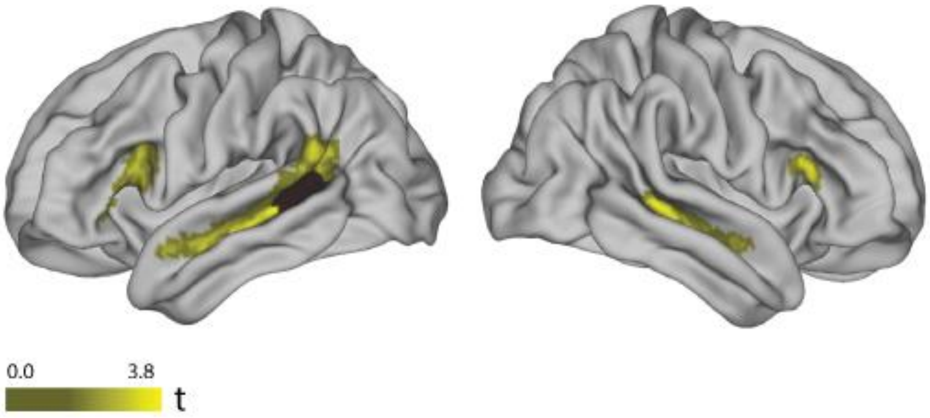


Figure S5. Positive parametric effect of total dependency length in sentences > word lists, Auditory sample, $P < .005$ uncorrected. In the LpMTG ROI, there was a significant Auditory > Visual effect, for sentences > word lists.

In discussion of these results, we note that our left-branching complexity measure, which targets overlapping non-adjacent dependencies, resulted in a more robust effect than the subtler effects observed using the (correlated) total dependency length measure. This pattern of results suggests that simultaneity (or overlap) of multiple unresolved non-adjacent dependencies, probes a partly different aspect of sentence processing than linear distance of non-adjacent dependencies (as indexed by the TDL measure). Both processes probably contribute to the difficulty of processing non-adjacent dependencies, but the simultaneity had the greater effect, at least on the BOLD-response. However, in most cases, these factors will be correlated, so the existing observations on difficulty associated linear distance (not controlling for simultaneity and direction of those dependencies) in the literature, are expected.

**Table S7**

Modality specific effects of total dependency length (TDL).

| Region | Cluster size | Cluster $p_{FWE\text{-}corr}$ | MNI- coordinates x | y | z | Voxel $p_{FWE\text{-}corr}$ | Voxel $T_{201}$ |
|---|---|---|---|---|---|---|---|
| TDL, Auditory > Visual<br>Sentences > word-lists | | | | | | | |
| *ROI LpMTG 10mm, SVC* | | | -50 | -42 | -4 | .036 | 2.99 |
| TDL, Auditory<br>Sentences > word-lists | | | | | | | |
| *LpMTG/STS* | 1281 | .021 | | | | | |
| *ROI LIFG, 10mm, SVC* | 121 | .025 | -48 | 16 | 18 | .010 | 3.44 |
| TDL, Auditory > Visual<br>Sentences | | | | | | | |
| *LSTG* | 1652 | .006 | | | | | |
| *RSTG* | 1518 | .009 | | | | | |
| TDL, Auditory<br>Sentences | | | | | | | |
| *LpMTG* | 3739 | .000 | | | | | |
| | | | -52 | -38 | 4 | .000 | 6.55 |
| | | | -54 | -12 | -4 | .002 | 5.32 |
| *RpMTG* | 2662 | .000 | | | | | |
| | | | 52 | -16 | -8 | .002 | 5.24 |
| | | | 50 | -34 | 2 | .009 | 4.90 |

No significant activations (neither at ROI-level) for: TDL, Sentences > words, Visual > Auditory; TDL, Sentences, Visual > Auditory
